# Soil microbial diversity and litter decomposition increase along a forest recovery gradient in tropical montane forests of Malaysian Borneo

**DOI:** 10.1101/2020.05.14.096883

**Authors:** Renee Sniegocki, Jessica B. Moon, Abigail L. Rutrough, Jude Gireneus, Jaya Seelan S. Seelan, David C. Weindorf, Michael C. Farmer, Kusum Naithani

**Affiliations:** University of Arkansas, Fayetteville, AR 72701 USA; Murray State University, Murray, KY 42071 USA; Texas Tech University, Lubbock, TX 79409 USA; Universiti Malaysia Sabah, Kota Kinabalu, Sabah, 88400, Malaysia

**Author notes:** Corresponding author: Kusum Naithani < > Address: SCEN 601, University of Arkansas, Fayetteville, AR, 72701, USA.

**Keywords:** 16s RNA, ITS, next-generation sequencing, soil microbiome, tea-bag index

## Abstract

Logging and forest conversion are occurring at alarming rates in the tropical forests. These disturbances alter soil chemistry and microbial diversity, and disrupt carbon cycling through shifts in litter decomposition. Direct links between microbial diversity and soil properties such as pH are well established; however, the indirect impacts of logging and forest conversion on microbial diversity and litter decomposition are poorly understood. We investigated how soil properties and soil functions change across a forest recovery gradient in the tropical montane forests of Malaysian Borneo. We used surface (top 5 cm) soil to assess soil physicochemical properties, next-generation DNA sequencing to assess soil microbial diversity, and standardized litterbags to assess litter decomposition and stabilization. Our results show that soils of the older forests harbored significantly greater microbial diversity, decomposed litter faster, and stabilized greater amounts of litter than soils of the younger forests and converted sites. These results suggest that logging and forest conversion significantly affect soil microbial diversity and can have lasting effects on carbon cycling in tropical montane forests.

## Introduction

Tropical forests are the most biodiverse ecosystems [1], store the largest amount of living carbon (C) [2], and account for over half of the global annual net primary production [3]. Logging and clear-cutting are major concerns in tropical forests, with an estimated 27.2 million ha cleared between 2000 and 2005 [4]. These activities can have lasting effects on canopy structure [5], microclimate [6], soil pH [7], soil nutrient content [8], C sequestration, productivity [9, 10], and multitrophic biodiversity [11, 12]. Southeast Asia is particularly vulnerable to this rapid loss of forests, where logged areas are often converted to more profitable rubber and oil palm plantations [13]. We investigated how logging and forest conversion affect soil properties and functions across a forest regeneration gradient in the tropical montane forests of Malaysian Borneo.

Disturbances such as logging and forest conversion can decrease soil C and nitrogen (N) content, pH, and a variety of other soil nutrients [7]. Disturbance-induced changes can also impact soil microbial communities because of the physical and chemical changes in soils [14–16]. While logging and forest conversion are capable of impacting microbial community composition and microbial biomass [17, 18], little is known about its role in shaping soil microbial diversity in the tropics. Microbes such as bacteria and fungi make up a significant portion of the biodiversity of tropical forests [19, 20]. While a number of studies found no difference in microbial diversity between logged and unlogged tropical forests [14, 21], others found an increase in microbial diversity along a tropical land-use intensification gradient [22, 23]. Despite the recent progress in our understanding of the soil microbiome, we still lack a clear understanding of how logging and forest conversion alter soil microbial diversity and functions in tropical ecosystems.

With an overarching goal of understanding how soil properties and functions change across a forest regeneration gradient in tropical montane forests, we ask: (1) *how do soil physical and chemical properties change along a forest regeneration gradient?* We expect to see changes in soil physical and chemical properties based on the number of years since the last major disturbance. Many prior studies have found disturbance events to significantly reduce soil pH and soil nutrients such as C and N [24, 25]. As an ecosystem recovers, these properties often return to near pre-disturbance levels [25, 26]. Given the consensus among prior studies, we hypothesize that soil physicochemical properties will show a directional change to recovery time. (2) *Does recovery time affect soil microbial diversity?* Microbial diversity has previously been shown to increase with recovery time in both tropical and temperate climates [27, 28]. Changes in fungal diversity are often attributed to successional changes in the plant community [26], while bacteria are often more strongly linked to soil properties such as pH [29]. We hypothesize that soil microbial diversity of tropical forests will increase with increasing recovery time. (3) *Do litter decomposition and stabilization change across the forest regeneration gradient?* We hypothesize that mature forests will have greater litter decomposition rates and litter stabilization compared to recently disturbed sites as reported by previous studies [30, 31].

## Materials and methods

### Study area

We selected five study sites along a forest regeneration gradient (time since the last major disturbance ranged from 4–150 year) in the Tambunan District of central Sabah, Malaysia (5.71750°, 116.40055°) (**Fig. 1** and **Table 1**). Two of the sites, the Mahua Falls Forest (MaF) and the Malungung Forest (MuF), were located within the federally protected Crocker Range Park area which covers ~ 139 919 ha. MaF and MuF were heavily logged ~100 and ~70 year (respectively) prior to our study. The remaining three sites were located in nearby privately-owned lands within the Tambunan Valley. The oldest site, Angelo’s Forest (AnF), had not been clear-cut/logged for over 150 year. The rubber site (Rub) was an abandoned rubber plantation cleared, terraced, and planted 41 year ago; trees at this site were untapped at the time of the sampling. The agricultural site (Agri) was previously cultivated for chili, cleared again in 2014, and left abandoned for four year prior to sampling. The study sites ranged in elevation from 870 to 1 150 m asl and experienced a similar regional climate with a mean annual temperature of 24.3 ℃ and annual precipitation of ~1 968 mm. The underlying geology of this region consists mainly of Quaternary fluvial gravels and sands [32], and the soils in this region are characterized as orthic acrisols, having low base saturation (< 50%) and high clay content in the B Horizon [33].

**Figure 1.**
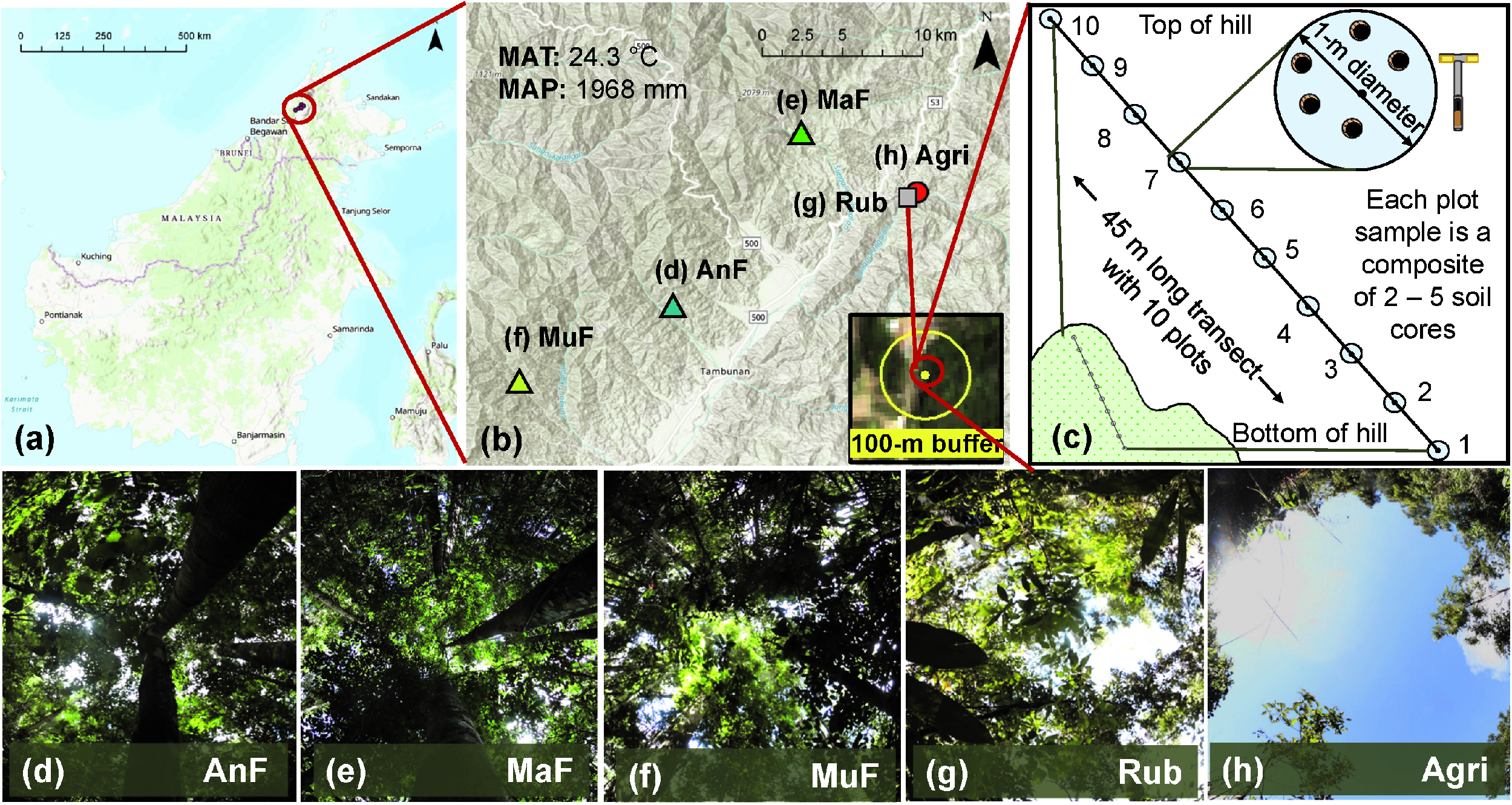
Description of the study sites and soil sampling scheme. Map of the study area (a) showing the location of the five study sites across a forest regeneration gradient in Sabah, Malaysian Borneo (b). The average elevation of the sites was 934 m, mean annual air temperature (MAT) was 24.3 ℃, and mean annual precipitation (MAP) was 1 968 mm. A 100-m buffer (yellow circle) was used around each site (yellow dot) to estimate percent forest cover using SENTINEL-2 imagery (b). Soil sampling scheme displaying a 45-m long transect with 10 sampling plots on an elevational gradient at each study site (c). Different sites including Angelo’s Forest (AnF), Mahua Falls Forest (MaF), Malungung Forest (MuF), Abandoned Rubber Plantation (Rub), and Abandoned Agriculture Field (Agri) are displayed using different colors and symbols (b) and corresponding canopy photos (d–h) are displayed below. Canopy photos were taken using a Canon EOS 7D Mark II Digital SLR camera with an Altura Ultra Wide Angle Aspherical Fisheye Lens (8 mm f/3.0).

**Table 1.** Study site description including geographical location (decimal degree), altitude (m +/- SE), slope (% +/- SE), dominant vegetative cover, recovery time (RT, year), and current/historical land use.

| Site Name | Latitude | Longitude | Altitude | Slope | Slope Aspect | Vegetation | RT | Land Use |
| --- | --- | --- | --- | --- | --- | --- | --- | --- |
| Angelo's Forest (AnF) | 5.713 | 116.335 | 900 ± 5 | 44 ± 19 | WNW | Mixed Dipterocarp forest | >150 | Not clear cut for >150 year; Currently forested. |
| Mahua Falls Forest (MaF) | 5.797 | 116.406 | 1140 ± 2 | 78 ± 0 | NNE | Mixed Dipterocarp forest | 100 | Clear cut ~100 year ago; Selective logging until approximately 1983. Currently forested. |
| Malungung Forest (MuF) | 5.661 | 116.251 | 862 ± 2 | 26 ± 4 | WNW | Mixed Dipterocarp forest | 70 | Clear cut ~70 year ago; Selective logging until 1993. Currently forested. |
| Rubber (Rub) | 5.765 | 116.469 | 879 ± 2 | 26 ± 4 | SSE | Rubber trees, roadside fern | 41 | Terraced and planted with rubber trees in 1977. Trees currently untapped and abandoned. |
| Agriculture (Agri) | 5.766 | 116.470 | 891 ± 1 | 56 ± 11 | SSW | Bamboo; mostly cleared | 4 | Previously cultivated for chili. Cleared in 2014. Currently used for agriculture. |

### Site Land Cover Classification

To describe current land use surrounding our sites, we classified land cover within a 100-m radius (**Fig. 1b**) of the transect center (**Fig. 1c**) using a maximum likelihood supervised classification. Maximum likelihood supervised classification uses pixel color and pattern recognition aided by user-provided training samples to classify satellite imagery into land cover classes [34]. We classified land cover as forest and non-forest in ArcMap (version 10.6.1, ESRI, Inc., © 1995–2018) using SENTINEL-2 satellite imagery. SENTINEL-2 imagery was downloaded from the Copernicus Open Access Hub (https://scihub.copernicus.eu/, last accessed 17 March 2019) and was selected due to its availability in our study area and high spatial resolution (10 m). Using all forested pixels within the 100-m radius buffer, we calculated percent forest cover for each site.

### Field Sampling

At each site, we collected the top 5 cm of soil (organic and mineral both) at 5-m intervals along a 45-m transect positioned along a slope gradient (**Fig. 1c, Table 1**), to sample maximum within-site variability, during the peak growing season of June 2018. We collected 2 – 5 soil cores (corer diameter = 6.35 cm) within 1 m of each plot center to create one composite soil sample per plot (**Fig. 1c**). Due to extremely rocky terrain at AnF, we were unable to use the 6.35 cm diameter soil auger and used a 2.54 cm diameter corer for soil sampling instead. We homogenized each composite sample and immediately preserved a 1 g subsample in RNALater stabilization solution (ThermoFisher Scientific). The preserved samples were shipped to the Molecular Research Laboratory (MR DNA, Shallowater, TX, USA) for microbial analysis within a week. In the remaining sample, we removed large roots, rocks, and undecayed litter. We then air-dried, ground (using pestle and mortar), and sieved (2 mm) the soil to remove non-soil fractions; and shipped these samples to the Texas Tech University’s soil laboratory for analysis of soil physicochemical properties. We measured surface (0 – 5 cm) soil temperature (digital soil thermometer probe, HANNA Instruments) in the field and collected additional surface soil samples in air-tight containers for assessing gravimetric soil water content (i.e., dried at 60 ℃ until sample weight remained constant). We measured air temperature and relative humidity every 30 min at the highest elevation along each transect using HOBO MX2301 data loggers (Onset Computer Corporation) throughout the growing season (**Fig. S1**).

### Soil microbial diversity

We received final operational taxonomic units (OTUs) that were assigned based on 97% similarity and followed the proprietary MR DNA analysis pipeline [35] from the Molecular Research Laboratory (MR DNA, Shallowater, TX, USA). See supplementary material for detailed information on molecular sample processing as provided by the lab. We calculated Chao1 Bacterial and Fungal diversity at the OTU level for each soil sample in R [36] using the package “SpadeR” [37].

### Soil physicochemical analysis

We measured soil pH using the saturated paste method [38], characterized soil elements (aluminum, sulphur, potassium, manganese, iron, nickel, copper, zinc, lead) using portable x-ray fluorescence (pXRF) [39], determined soil organic matter (SOM, %) concentration using the loss-on-ignition method [40], estimated total C and N using the Dumas method [41], and determined soil texture (i.e., % sand, silt, clay) using a Hydrometer particle size analysis (152H Hydrometer) [42]. Due to insufficient sample size, we performed soil texture analysis on 28 soil samples out of 50 but the data provided a general idea of the soil type at each site (**Fig. S2**).

### Litter decomposition and stabilization

Inherent differences in litter quality can affect microbial decomposition rates following disturbance [43]. The Tea Bag Index method employs a standardized litter bag technique to compare plant litter decomposition and stabilization across sites without the influence of litter quality; and focuses on identifying environmental factors that influence decomposition rates across sampling locations [43]. We buried 10 pairs of pre-weighed litter bags (i.e, nonwoven tetrahedron-shaped polypropylene tea bags, Lipton, Unilever), one green tea (fast decomposing litter) and one rooibos tea (slow decomposing litter), at 8 cm below surface. After 72 to 77 days, we retrieved the teabags and manually removed debris and roots following the “Tea Bag Index: Scientific Protocol, 2016” (http://www.teatime4science.org/method/stepwise-protocol/). We used the initial and final oven-dried (70 ℃ for 48 hours) mass to calculate the decomposition rate constant (*k*, d^−1^) and stabilization factor (*S*, unitless).

### Statistical Analyses

First, we used SigmaPlot 14.0 (Systat Software Inc., San Jose California, USA) to display and fit the statistical models to the bivariate relationships between recovery time and different soil properties. All regression models were evaluated for strength (simpler models with greater R^2^ were selected) and significance (only significant (p ≤ 0.05) models were selected) (**Fig. 3**).

Then, we used the “randomForest” package [45] in R to evaluate the relative importance of recovery time as a predictor of soil microbial diversity and soil functions (litter decomposition and stabilization) when compared with other predictors including soil physicochemical properties, aboveground vegetation cover, and environmental variables. Random forest is an ensemble machine learning technique that creates a series of uncorrelated regression trees from random subsamples of the data. These trees decide as a committee which variables are the most predictive, and rank the variables in order of their ability to increase the mean square error when removed from the model [46]. Before running the random forest analysis, we used the package “corrplot” in R to identify and remove highly correlated variables (|r| > 0.70) [44] (**Fig. 2**), retaining the most ecologically relevant and/or proximal predictor variables [47]. Through this process iron, C, manganese, sulfur, potassium, carbon:N, and zinc were removed from the random forest analysis. Although recovery time and pH were highly correlated (|r| = 0.82), we chose to leave both variables in the analysis because of their unique ecological contributions to the model. To ensure that the importance given to each predictive variable would not be biased by the presence of the two correlated variables, we ran the random forest analyses again removing each of the two correlated variables one at a time and then simultaneously. We found reductions in the overall variance explained by the model, however the top predictive variables for each model were the same, excluding the variable(s) removed. Therefore, we concluded that the correlation of the variables had little effect on our model and chose to include them both in the final analyses. One extreme fungal diversity outlier was removed from the random forest and regression analyses, but is displayed in figures and noted with an asterisk. We built 200 trees (ntrees = 200) for each run and sampled 3 – 4 variables (Mtry = 3 – 4, depending on the number of predictor variables) with replacement for each tree. Finally, we took the top four predictors and used regression analysis to further explore the trends (**Fig. 4 – 7)**. Regression models of bivariate relationships were fitted using SigmaPlot 14.0.

**Figure 2.**
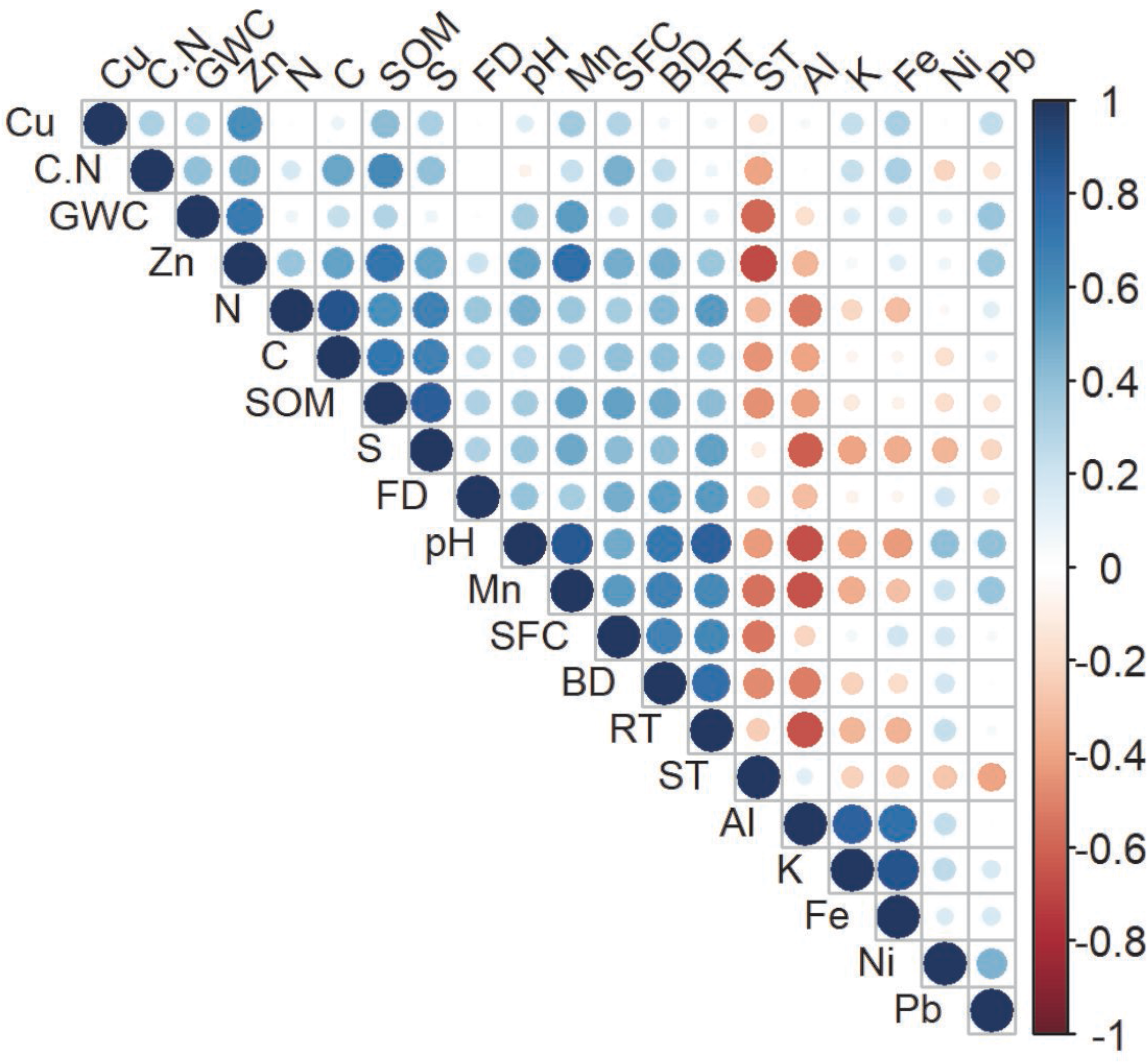
Correlation matrix of measured variable including recovery time (RT, year), surrounding forest cover (SFC, %), daytime soil temperature (ST, °C), gravimetric water content (GWC, g g^−1^), pH, soil organic matter (SOM, %), carbon:nitrogen ratio (C.N), Cu (mg g^−1^), Zn (mg g^−1^), N (%); C (%), S (mg g^−1^), Mn (mg g^−1^), Al (mg g^−1^), K (mg g^−1^), Fe (mg g^−1^), Ni (mg g^−1^), Pb (mg g^−1^), and soil microbial diversity (BD: bacterial OTU Chao1 diversity index and FD: fungal OTU Chao1 diversity index).

**Figure 3.**
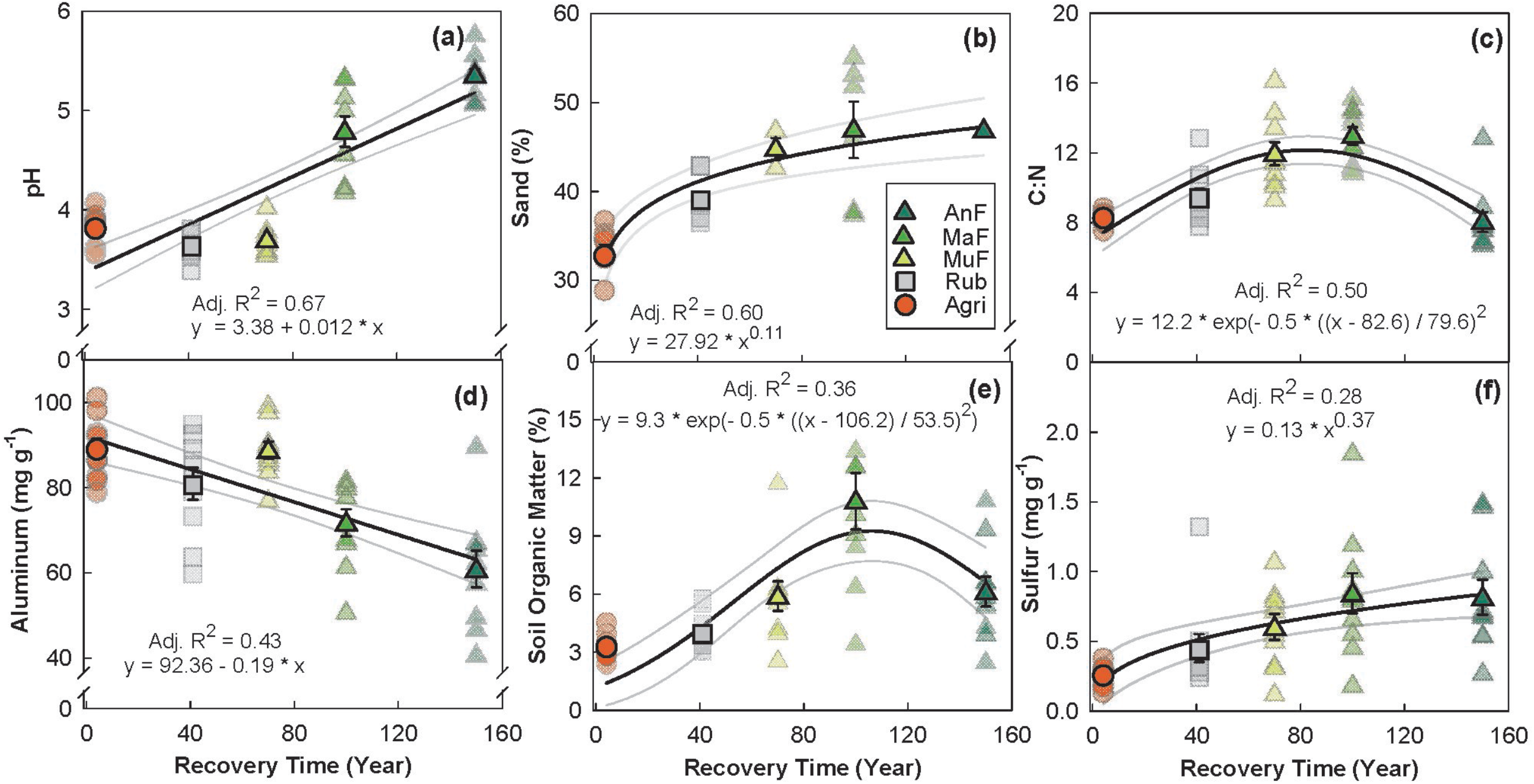
Relationship between recovery time (year) and soil physicochemical properties (a – f) across a forest regeneration gradient: Angelo’s Forest (AnF), Mahua Falls Forest (MaF), Malungung Forest (MuF), Abandoned Rubber Plantation (Rub), and Abandoned Agriculture Field (Agri). Smaller symbols in the background of each plot represent raw data points (n = 10), while larger symbols represent site means. Sample size for for % sand varied from 1-10 depending on the site (AnF (n=1), MaF (n=6), MuF (n=3), Rub (n=8), Agri (n=10)) due to logistical difficulties of collecting enough soil with the 5 cores. Bars represent the standard error of mean and gray lines represent 95% confidence intervals. All statistical models are significant at p ≤ 0.05.

**Figure 4.**
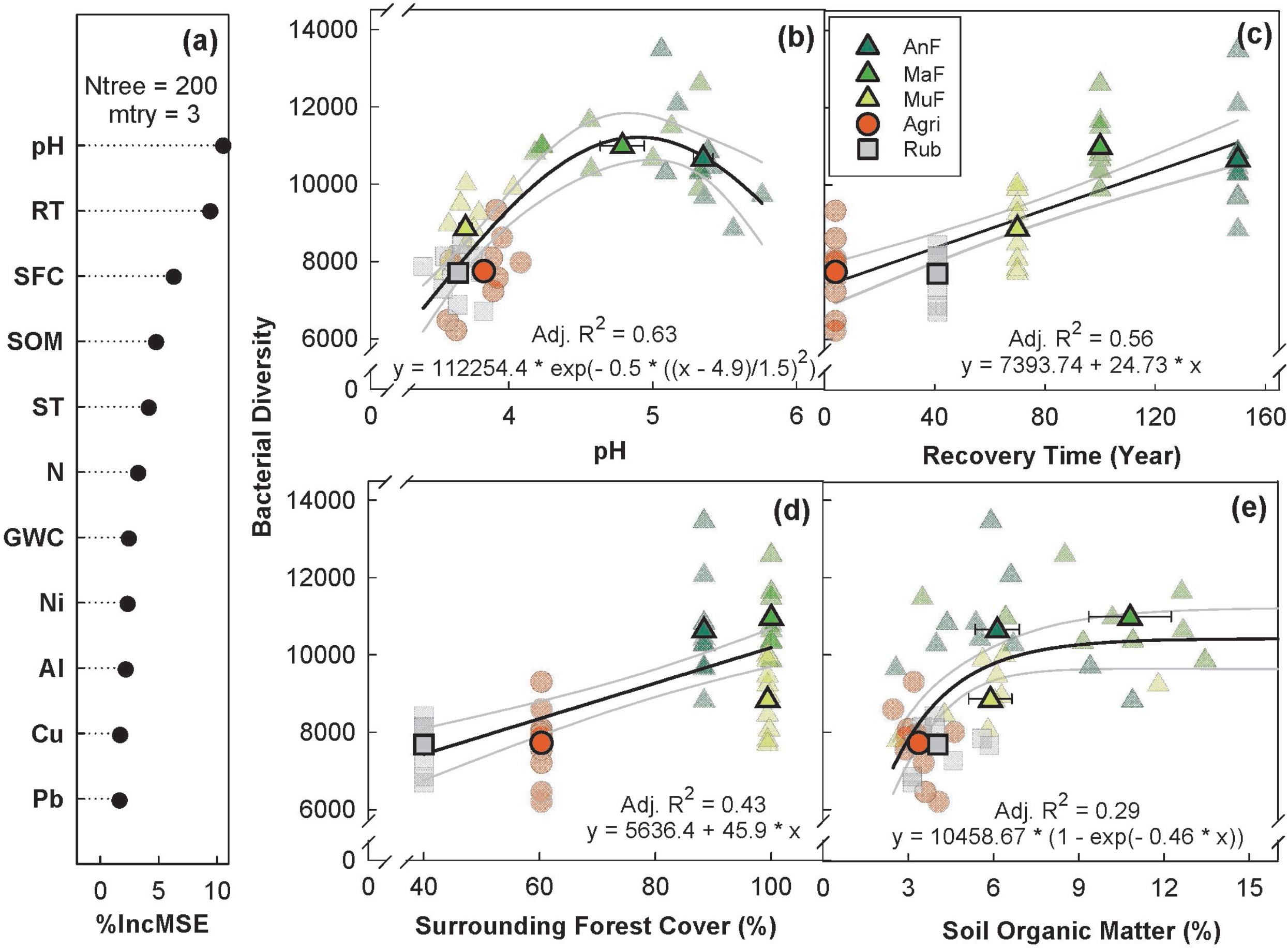
Variable importance prediction from random forest analysis (a) and bivariate relationships of top four predictors (b – e) of bacterial diversity (Chao1 diversity index) across a forest regeneration gradient. Soil variables included in random forest analyses, in order of importance, are: pH, recovery time (RT, year), surrounding forest cover (SFC, %), soil organic matter (SOM, %), daytime soil temperature (ST, °C), N (%), gravimetric water content (GWC, g g^−1^), Ni (mg g^−1^), Al (mg g^−1^), Cu (mg g^−1^), and Pb (mg g^−1^). Different sites including Angelo’s Forest (AnF), Mahua Falls Forest (MaF), Malungung Forest (MuF), Abandoned Rubber Plantation (Rub), and Abandoned Agriculture Field (Agri) are displayed using different colors and symbols. Smaller symbols in the background of each bivariate plot represent raw data points (n = 10), while larger symbols represent site means. Bars represent the standard error of mean and gray lines represent 95% confidence intervals. All statistical models are significant at p ≤ 0.05.

## Results

### Surrounding forest cover

Surrounding forest cover varied among the study sites, with 100% forest cover measured at MaF, 99.4% at MuF, 88.4% at AnF, 60.3% at Agri, and 40% at Rub. While surrounding forest cover was not significantly related to recovery time, higher surrounding forest cover was found at the three naturally forested sites than the agricultural and converted sites (**Fig. S3**).

### Effect of recovery time on soil physicochemical properties

Soil pH (p < 0.001, Adj R^2^ = 0.67), sulfur (p = 0.05, Adj. R^2^ = 0.28) and % sand (p < 0.001, Adj. R^2^ = 0.60) increased linearly while aluminum decreased linearly (p < 0.001, Adj. R^2^ = 0.42) with increasing recovery time. SOM (p < 0.001, Adj. R^2^ = 0.36) showed a peaked response at ~100 year following recovery (**Fig. 3**). Total N content increased linearly (p < 0.001, Adj. R^2^ = 0.31) while total C content (p < 0.001, Adj. R^2^ = 30) showed a saturated response with recovery time; taken together this generated a peaked response of C:N at ~100 year following recovery (**Fig. 3, Fig. S3**).

### Relative influence of recovery time on soil microbial diversity

#### Bacterial diversity

Random forest analysis explained 67.08% of the total variance in Chao1 bacterial OTU diversity (hereafter, bacterial diversity) with pH, recovery time, percent forest cover, and SOM as top four predictors (**Fig. 4a–e**). Bacterial diversity peaked at a pH of 5 (p < 0.001, Adj. R^2^ = 0.63, **Fig. 4b**), and increased with both recovery time (p ≤ 0.01, Adj. R^2^ = 0.56, **Fig. 4c**), and surrounding forest cover (p < 0.001, Adj. R^2^ = 0.43, **Fig. 4d**). Bacterial diversity levels also saturated around 6% SOM (p < 0.001, Adj. R^2^ = 0.29, **Fig. 4e**).

#### Fungal diversity

Random forest analysis explained 18.69% of the total variance in Chao1 fungal OTU diversity (hereafter, fungal diversity); recovery time, percent forest cover, pH, and Al were the top four predictors (**Fig. 5a–e**). Fungal diversity increased with recovery time saturating at ~100 year (p < 0.001, Adj. R^2^ = 0.34, **Fig. 5b**). Surrounding forest cover was the second most important predictor of fungal diversity, with a positive linear relationship (p < 0.001, Adj. R^2^ = 0.21, **Fig. 5c**). Fungal diversity showed a peaked response to pH with peak at ~ 5 (p ≤ 0.01, Adj. R^2^ = 0.17, **Fig. 5d**) and a weak inverse relationship with soil aluminum concentration (p ≤ 0.01, Adj. R^2^ = 0.09, **Fig. 5e**).

**Figure 5.**
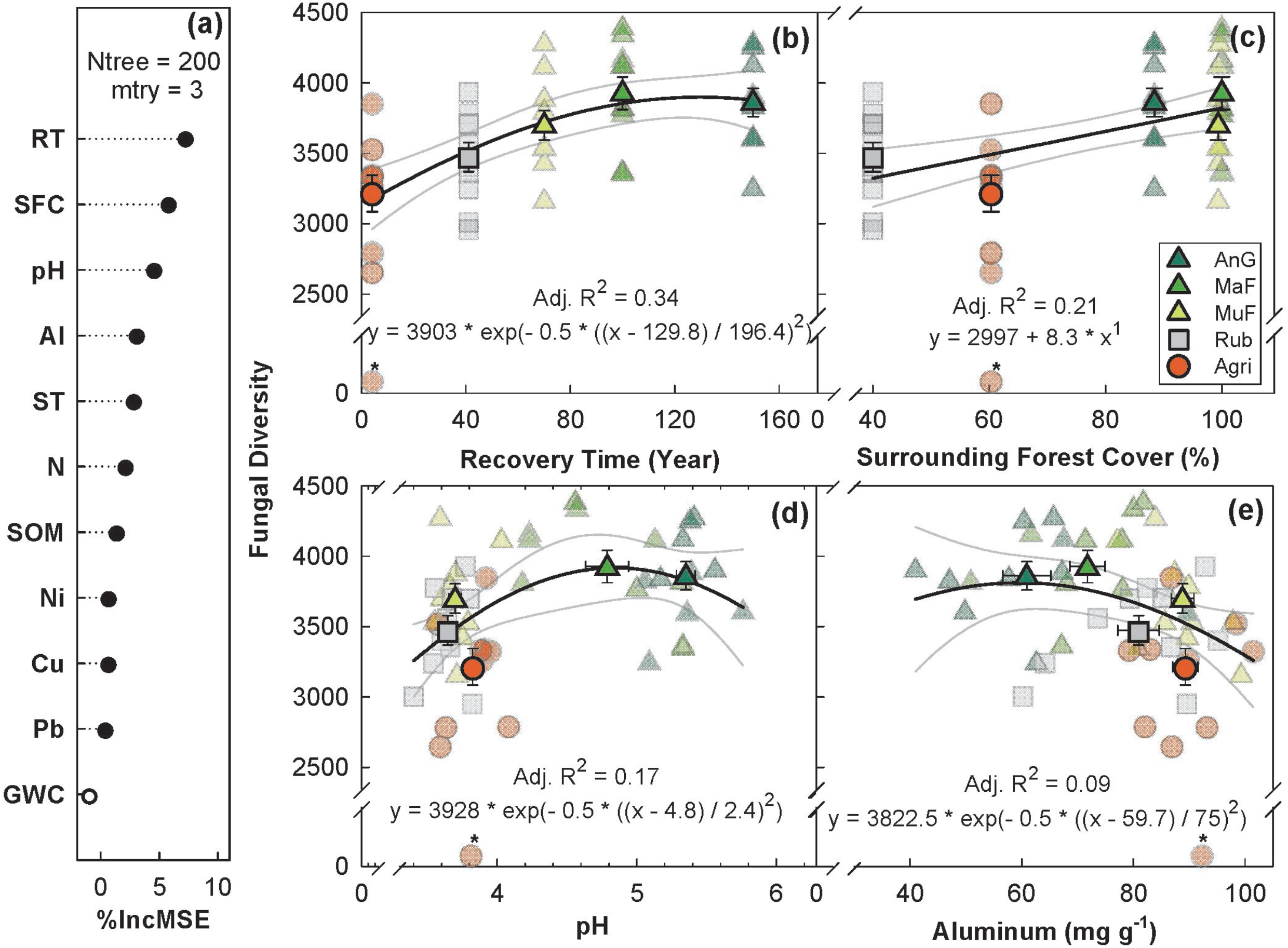
Variable importance prediction from random forest analysis (a) and bivariate relationships of top four predictors of fungal diversity (Chao1 diversity index) across a forest regeneration gradient (b – e). Soil variables included in random forest analyses, in order of importance, are: recovery time (RT, year), surrounding forest cover (SFC, %), pH, Al (ppm), daytime soil temperature (ST, °C), N (%), soil organic matter (SOM, %), Ni (mg g^−1^), Cu (mg g^−1^), Pb (mg g^−1^), gravimetric water content (GWC, g g^−1^, not significant). Different sites including Angelo’s Forest (AnF), Mahua Falls Forest (MaF), Malungung Forest (MuF), Abandoned Rubber Plantation (Rub), and Abandoned Agriculture Field (Agri) are displayed using different colors and symbols. Smaller symbols in the background of each bivariate plot represent raw data points (n = 10), while larger symbols represent site means. Bars represent the standard error of mean and gray lines represent 95% confidence intervals. All statistical models are significant at p ≤ 0.05.

### Relative influence of recovery time on litter decomposition and stabilization

#### Litter decomposition rate constant (*k*)

Random forest analysis explained 15.08% overall variance in *k* with N, pH, fungal diversity, and bacterial diversity as the top four predictors (**Fig. 6a–e**). *k* increased linearly with increasing soil N (p < 0.05, Adj. R^2^ = 0.22, **Fig. 6b**), fungal (p ≤ 0.05, Adj. R^2^ = 0.22, **Fig. 6d**) and bacterial diversity (p < 0.001, Adj. R^2^ = 0.19, **Fig. 6e**) and peaked at a soil pH of ~ 5 (p < 0.05, Adj. R^2^ = 0.13, **Fig. 6c**). *k* increased with recovery time, saturating at ~100 year (p < 0.05, Adj. R^2^ = 0.15, **Fig. S3**). Our *k* estimates are comparable to other lowland tropical forests from a global dataset (**Fig. S4)**, with relatively high *k* estimates in the two most mature forests (0.0320 ± 0.005 (AnF) and 0.0357 ± 0.006 (MaF) d^−1^).

**Figure 6.**
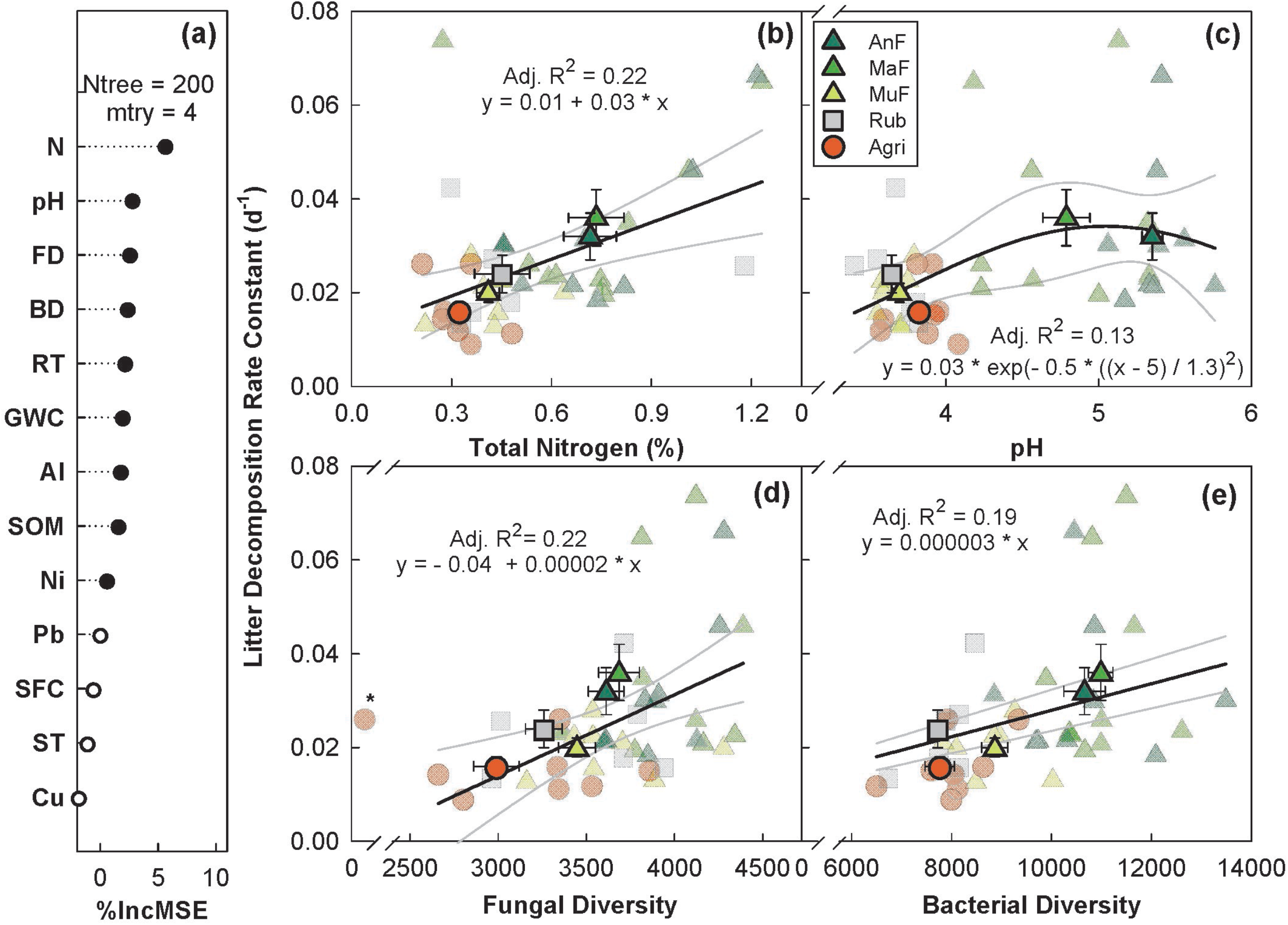
Variable importance prediction from random forest analysis (a) and bivariate relationships of top four predictors of litter decomposition rate across a forest regeneration gradient (b – e). Significant (filled circles) soil variables included in random forest analyses, in order of importance, are: N (%), pH, fungal diversity (FD, Chao1 diversity index), bacterial diversity (BD, Chao1 diversity index), recovery time (RT, year), gravimetric water content (GWC, g g^−1^), Al (mg g^−1^), soil organic matter (SOM, %), and Ni (mg g^−1^). Different sites including Angelo’s Forest (AnF), Mahua Falls Forest (MaF), Malungung Forest (MuF), Abandoned Rubber Plantation (Rub), and Abandoned Agriculture Field (Agri) are displayed using different colors and symbols. Smaller symbols in the background of each bivariate plot represent raw data points (n = 10), while larger symbols represent site means. Bars represent the standard error of mean and gray lines represent 95% confidence intervals. All statistical models are significant at p ≤ 0.05.

#### Litter stabilization factor (*S*)

Random forest analysis explained 47.09% of the overall variance in *S* with surrounding forest cover, bacterial diversity, recovery time, and daytime soil temperature as the top four predictors (**Fig. 7a**). *S* increased with surrounding forest cover, saturating at ~ 80% cover (p < 0.005, Adj. R^2^ = 0.63, **Fig. 7b**). *S* peaked at mid-range bacterial diversity (p < 0.001, Adj. R^2^ = 0.32, **Fig. 7c**), and increased linearly with recovery time (p < 0.01, Adj. R^2^ = 0.14, **Fig. 7d**). *S* also increased with daytime soil temperature, but peaked between 20 - 21 ℃ (p < 0.001, Adj. R^2^ = 0.22, **Fig. 7e**). All three naturally regenerating forests had higher *S* estimates (AnF = 0.239 ± 0.014, MaF = 0.245 ± 0.011, and MuF = 0.242 ± 0.017) than Rub (*S* = 0.070 ± 0.025) and Agri (*S* = 0.201 ± 0.017) sites.

**Figure 7.**
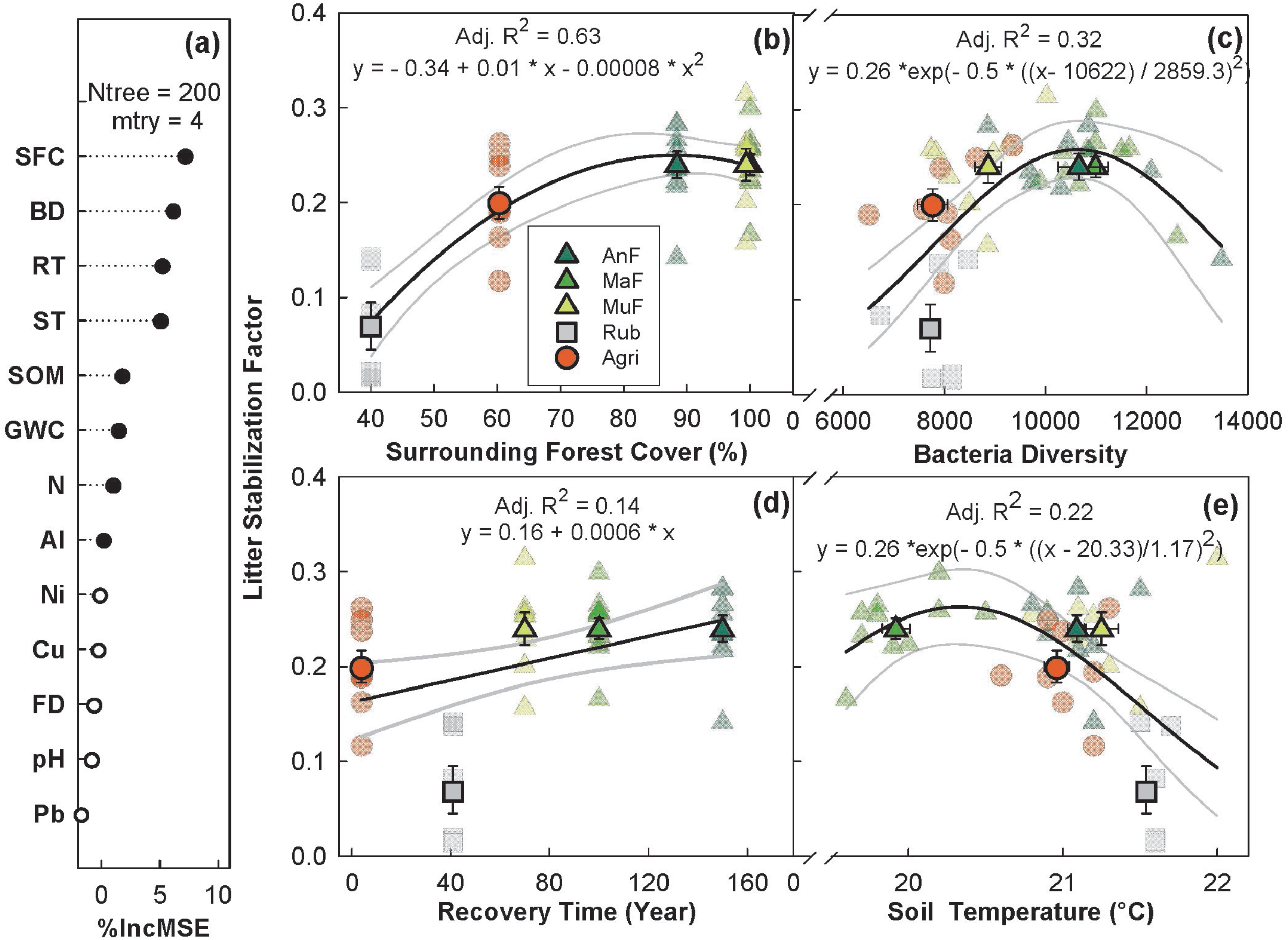
Variable importance prediction from random forest analysis (a) and bivariate relationships of top four predictors of litter stabilization across a forest regeneration gradient (b – e). Significant (filled circles) soil variables included in random forest analysis, in order of importance, are: surrounding forest cover (SFC, %), bacterial diversity (BD, Chao1 diversity index), recovery time (RT, year), daytime soil temperature (ST, °C), soil organic matter (SOM, %), GWC (gravimetric water content, g g^−1^), Ni (mg g^−1^), and Al (mg g^−1^). Different sites including Angelo’s Forest (AnF), Mahua Falls Forest (MaF), Malungung Forest (MuF), Abandoned Rubber Plantation (Rub), and Abandoned Agriculture Field (Agri) are displayed using different colors and symbols. Smaller symbols in the background of each bivariate plot represent raw data points (n = 10), while larger symbols represent site means. Bars represent the standard error of mean and gray lines represent 95% confidence intervals. All statistical models are significant at p ≤ 0.05.

## Discussion

Logging and forest conversion continue to craft the tropical montane landscape, with limited understanding of the consequences to global C cycling. In this study, we show strong links between forest regeneration and soil physicochemical properties, microbial diversity, and C cycling in a tropical montane region of Sabah, Borneo. We found that many of the physical and biological variables covary across this forest regeneration gradient and take ~100 year to reach pre-disturbance levels; this highlights the vulnerability of these systems to the effects of agroforestry use.

### Effect of recovery time on soil physicochemical properties

We compared soil properties at five sites across a forest regeneration gradient, with sites ranging from 4 to over 150 year of post-disturbance regeneration. As expected, recovery time significantly affected soil physicochemical properties (**Fig. 3**). Soil pH had the most significant change along the gradient, increasing from 3.7 at the youngest site (Agri) to 5.4 at the oldest site (AnF) **(Fig. 3a)**. This pH range is similar to the range reported by others [21] across a forest disturbance gradient in Sabah, Borneo, and consistent with findings in other tropical systems [21, 24, 25] that reported lower pH after forest logging and conversion. Lower pH levels measured in recently recovered sites could be attributed to recent harvesting. For example, plants are capable of altering the pH of the rhizosphere and bulk soil through ion uptake, and removal of soil cations such as Ca^2+^ and Mg^2+^ can lead to soil acidification when plant material is removed from a site during harvest [48]. Previous logging and cultivation of the sites may have resulted in a decrease in soil pH, with soil pH gradually increasing over time in older forests with organic matter decomposition [49].

We found that sulfur increased, while SOM, C content, and C:N showed peak or saturating responses with recovery time (**Fig. 3**). Our results are consistent with prior work that reported higher concentrations of total soil C, N [50], and SOM [25] in forested sites compared to both agricultural and plantation sites, mainly attributed to a decrease in quantity and quality of plant residues returned to the soil. Though there is no information on how sulfur changes along a forest regeneration gradient, soil sulfur concentrations can be tightly coupled with total soil C content and SOM levels [51]. Our results reflect this nutrient coupling, with the soil sulfur concentration increasing proportionally with soil C across the forest regeneration gradient. Previous studies have observed significant losses of soil nutrients due to leaching [52], biomass removal [53, 54], and altered nutrient allocation [55] associated with logging. The increased soil C pools and nutrients along this forest regeneration gradient are possibly due to the soil nutrient stocks rebuilding to pre-disturbance levels.

Given that rates of organic matter accumulation typically slow over time [56], the observed peaked relationships were not surprising. Percent SOM has been found to return to pre-disturbance levels within 40–50 year of succession in tropical forests [57], however, other studies have observed this to occur in less than a decade [25]. The recovery time required to accumulate organic matter in the soil is variable and can be affected by vegetation (i.e., quantity and quality of litter inputs), abiotic and biotic soil properties (i.e., soil texture and microbial activity), and land use history [58]. Furthermore, inherent differences among our forested sites in terms of soil texture and terrain (the oldest site, AnF, had rocky terrain) could have resulted in the high variability and lower amounts of SOM as compared to the other old growth forest (MaF). Similar to soil C and SOM, the C:N peaked at ~100 year and subsequently declined at the oldest site, due to the combined effects of total soil C decreasing and total soil N increasing (**Fig. 3 and Fig. S3**). Nitrogen fixing organisms, particularly microbial plant symbionts, are considered an important source of new N in tropical soils [59]. Some late-successional plant species have higher leaf N content, with correspondingly high soil N content, as compared with early-successional forests [60]. Greater N content in litter could result in enhanced decomposition rates, driving soil C and SOM levels down.

Soil texture also changed significantly along the recovery gradient; % sand increased until around 100 year post-recovery (**Fig. 3**). Fine particles can be lost from soil following disturbance events such as tillage due to erosion and runoff, leaving a higher percentage of larger particles such as sand [61]. Our results contradict these findings and could be a result of a number of factors including variations in soil parent material and natural erosion patterns. While we identified no effects of recovery time on soil microclimate, we found a decrease in daytime soil temperature with increasing forest cover (**Fig. S5**). Removal of the forest canopy during logging events increases daytime soil temperature [62], and reforestation can have significant surface cooling effects [63].

### Relative influence of recovery time on soil microbial diversity

#### Bacterial diversity

Our results show that soil pH is the strongest overall predictor of bacterial diversity (**Fig. 4b**), which is consistent with previous work on bacterial diversity in the tropics [21, 29], temperate forests, grasslands [64–66] and alpine meadows [67]. Bacterial diversity peaked at a pH of ~ 5; this is consistent in pattern with prior work on forested systems [29, 68], but peak diversity occurred at a pH of ~ 7 in one of these studies [29]. Bacteria are known to experience physiological constraints at extreme pH levels and optimum soil pH varies among bacterial taxa [29]. However, the intracellular pH of bacteria is often close to neutral [69, 70], and a more extreme external pH may place stress on bacterial cells [71]. The current understanding of the relationship between bacterial diversity and pH suggests that changes in bacterial diversity may be the result of pH-driven changes in availability of various soil nutrients [29, 71].

Our results showed that bacterial diversity increased linearly with recovery time (**Fig. 4c**), which is consistent with prior studies [27, 28] that found an increase in microbial diversity with stand age. While previous work has reported a peak in bacterial diversity at ~ 40 year post-disturbance in a subtropical secondary forest [28], our study showed no decline in bacterial diversity. We found that the surrounding forest cover is a significant predictor of bacterial diversity (**Fig. 4d**). The lowest bacterial diversity was found in the landscapes dominated by human activities (Rub and Agri), evident from the existence of social trails, grazing (Agri), amount of exposed soil, and the proximity to inhabited areas. The three naturally regenerated forested sites (MuF, MaF, and AnF) were relatively undisturbed and surrounded by a large tropical forest buffer. Previous research suggests that anthropogenic disturbance such as conversion to pasture or plantation can induce biotic homogenization [72, 73] of the soil bacterial community, leading to the loss of some more endemic taxa and to the broader dispersal of the existing taxa within a site [24, 74]. This pattern is also supported by our findings that the naturally forested sites showed the most variation along the transect for a number of soil properties, including pH, SOM, and total C content. Associated losses of endemic taxa are likely to be better quantified by the Chao1 diversity index, used in this study, as it is weighted by rare taxa. SOM was the fourth most important predictor of bacterial diversity, showing a saturating response of bacterial diversity at ~ 6% SOM (**Fig. 4e**). Soil bacterial communities rely on the decomposition of organic matter for energy [75], and SOM has previously been identified as one of the main drivers of soil bacterial diversity on a global scale [76].

#### Fungal diversity

We found that fungal diversity increased significantly with recovery time until ~100 year (**Fig. 5b**), which is consistent with prior studies [27, 28] that found an increase in microbial diversity with stand age. This increase in microbial diversity over time may reflect spatial heterogeneity in accumulation of SOM [77] or increased variability in litter quality and quantity as a result of changes in plant diversity [28] as forests mature. Soil disturbance and homogenization from tillage and logging can negatively impact fungal communities [78]. As mentioned earlier, disturbed sites, Agri and Rub, showed lower variation along the transect for a number of soil parameters including pH, SOM, and total C content. These findings suggest that the sites incurred resource homogenization during forest clearing, cultivation, and continued anthropogenic disturbance. Many fungal species (e.g., namely ectomycorrhizal fungi), are plant-symbionts and often exhibit strong host tree preference [79]. Thus, the alteration of tree species composition and abundance can also have direct effects upon fungal community structure and diversity [80]. We found that the current surrounding forest cover (i.e., cover by a mature tree canopy) is a significant predictor of fungal diversity, with Agri and Rub exhibiting the lowest levels of fungal diversity, and mature forest exhibiting higher fungal diversity (**Fig. 5c**). Future studies in this region, relating plant community cover, composition, and root structure to the relative abundance of fungal taxa, would aid in establishing these linkages.

Overall, random forest analysis explained only 18.69% of variation in fungal diversity. The lack of strong direct ties between fungal diversity and soil physicochemical properties could be tied to the fungal scale of operation; fungal hyphae can access a larger volume of soil, making fungi more able to survive in a variety of soil conditions, while bacteria are more heavily impacted by local substrate and nutrient availability [81]. Of the physicochemical properties evaluated we found that soil pH and soil Al concentrations, which were negatively correlated with each other (|r| = 0.66), were the strongest predictors of fungal diversity (**Fig. 5d, e**). Although similar results have been found in previous studies [81] these relationships are often weaker [82, 83] than those between soil pH and bacterial community composition and diversity [21, 29, 66, 71].

### Relative influence of recovery time on litter decomposition and stabilization

#### Litter decomposition rate (*k*)

While recovery time isn’t a strong predictor of *k*, microbial diversity and some soil physicochemical properties are strongly correlated with *k*. Our results suggest that soil N is the most important predictor of *k* (**Fig. 6b**) followed by pH (**Fig. 6c**) and microbial diversity (**Fig. 6d–e**). Because microbially mediated litter decomposition is generally thought to be nutrient limited [87], it is likely that increases in nutrient levels in these ecosystems increased *k*. These results are consistent with a previous meta-analysis [88] that found that N addition often increased decomposition of high-quality litter. Soil microbial communities are one of the main drivers of soil C cycling through processes such as litter decomposition and stabilization [89]. Previous studies, focused on the relationship between microbial diversity and soil functions, have found mixed results with some showing little to no effect of microbial diversity on litter decomposition [90] and others finding positive relationships between the two [91]. We found that *k* increased linearly with increasing bacterial (**Fig. 6d**) and fungal (**Fig. 6e**) diversity, suggesting a broader metric such as diversity can be used as an indicator of functional shifts in the tropical montane systems. These findings also have significant implications on current debate about diversity versus functional redundancy [92], by suggesting that reduced microbial diversity results in reduced functional capabilities of the soil.

#### Litter stabilization factor (*S*)

*S* increased linearly with recovery time and showed a saturating response to surrounding forest cover, reaching a maximum at ~ 90% cover in the older forests (AnF and MaF, **Fig. 7**). Possible mechanisms for this stabilization include the formation of a specialized microbial community with distinct stabilizing capabilities [93], the addition of a more preferential microbial energy source through root exudates [94], the increased input of recalcitrant root tissue, the physical protection of organic matter by the formation of soil aggregates [95], and reduced erosion and leaching [30]. During litter decomposition, some labile plant tissues are microbially and biochemically transformed into recalcitrant, stable compounds. The diversion of labile plant tissues, which would otherwise decompose, into stabilized organic matter builds upon humus pools, thereby sequestering more C in the soil [95]. Previous studies have found that the presence/absence of woody vegetation [31] can affect C turnover. It is likely that increased forest cover and live root mass protected the litter bags from decomposition and led to higher stabilization. The three forested sites (AnF, MaF, and MuF) had the highest *S* followed by Agri, and Rub had the lowest stabilization among the sites. The youngest site (Agri) had relatively low tree cover, but had a dense herbaceous understory that could have provided some physical protection to the soil, an effect that has been previously documented in fallow land in tropical montane regions [30]. Rub site on the other hand, had a considerable amount of exposed soil both beneath the tree cover and in the surrounding area.

Litter stabilization showed no relationship with fungal diversity (**Fig. 7a**), but a peaked relationship with bacterial diversity, with peak at mid-range of bacterial diversity (**Fig. 7c**). Changes in microbial growth efficiency, or the ability of the microbial community to incorporate substrates into biomass and byproducts, can alter litter transformation rates [96]. Our data matched the global patterns of litter decomposition and stabilization, but both processes were decoupled unlike the global data which showed an inverse relationship between *k* and *S* [43] (**Fig. S4**).

Considering the rapid rate of deforestation in the tropics and microbial role in nutrient cycling, it is critical to investigate how logging and forest conversion affect soil structure and functions in these systems. Our results suggest that logging and forest conversion significantly affect soil microbial diversity and can have lasting effects on C cycling in tropical montane forests. Considering the potential global impacts of land-use on the terrestrial C cycle in one of the most productive biomes on Earth, these results have wide implications for land management and biodiversity conservation.

## Author Contribution

KN designed the study; MF collaborated with private landowners for land access and selected sites across a gradient of land use disturbance; RS, JBM, AR, JG, and KN conducted field work; DCW conducted soil physicochemical analysis; RS analysed the data with assistance from KN; RS wrote the manuscript with assistance from KN, JBM, and JSSS; all authors approved the paper.

## Acknowledgements

The NSF IRES grants (1658711, 1658722) funded this study and the data was collected under the permit (# JKM/MBS.1000 - 2/3 JLD.3(41)) from the Sabah Biodiversity Council. We thank Angelo Dosis Asis, Jerry Peter Wala, and Walter Sorop for providing access to their land and helping with logistics. We thank Drs. Tigga Kingston, Robin Verble, and Sarah Fritz for helping with the travel and logistics of IRES related activities, and Joumin Rangkasan for assistance in the field. DCW gratefully acknowledges the BL Allen Endowment in Pedology at Texas Tech University in conducting this research.

