## Supplementary Material for "Soil microbial diversity and litter decomposition increase along a forest recovery gradient in tropical montane forests of Malaysian Borneo"

### **Content**

1. Tables
   a. Table S1
2. Figures
   a. Figure S1
   b. Figure S2
   c. Figure S3
   d. Figure S4
   e. Figure S5
3. Methods
   a. Soil Microbial analysis

**Table S1:** Soil properties from the five study sites, including pH in water, gravimetric water content (g g^-1^), daytime soil temperature (℃), soil organic matter (%), and carbon to nitrogen ratios (C:N). Sites are arranged from longest to shortest time since disturbance.

| **Site Name** | **pH** | **Soil Water Content** | **Soil Temp** | **Soil Organic Matter** | **C:N** |
| --- | --- | --- | --- | --- | --- |
| Angelo's Forest (AnF) | 5.35 ± 0.07 | 0.47 ± 0.04 | 21.09 ± 0.06 | 6.13 ± 0.78 | 8.04 ± 0.57 |
| Mahua Falls Forest (MaF) | 4.79 ± 0.15 | 1.01 ± 0.16 | 19.92 ± 0.09 | 10.80 ± 1.45 | 12.97 ± 0.50 |
| Malungung Forest (MuF) | 3.70 ± 0.04 | 0.40 ± 0.11 | 21.25 ± 0.11 | 5.89 ± 0.76 | 11.94 ± 0.67 |
| Rubber (Rub) | 3.65 ± 0.04 | 0.54 ± 0.03 | 21.54 ± 0.03 | 4.03 ± 0.31 | 9.44 ± 0.49 |
| Agriculture  (Agri) | 3.82 ± 0.05 | 0.51 ± 0.08 | 20.96 ± 0.08 | 3.36 ± 0.20 | 8.32 ± 0.12 |

**Figure S1.** Air temperature (℃) and relative humidity (%) measurements every 30-min interval for the duration of this study.

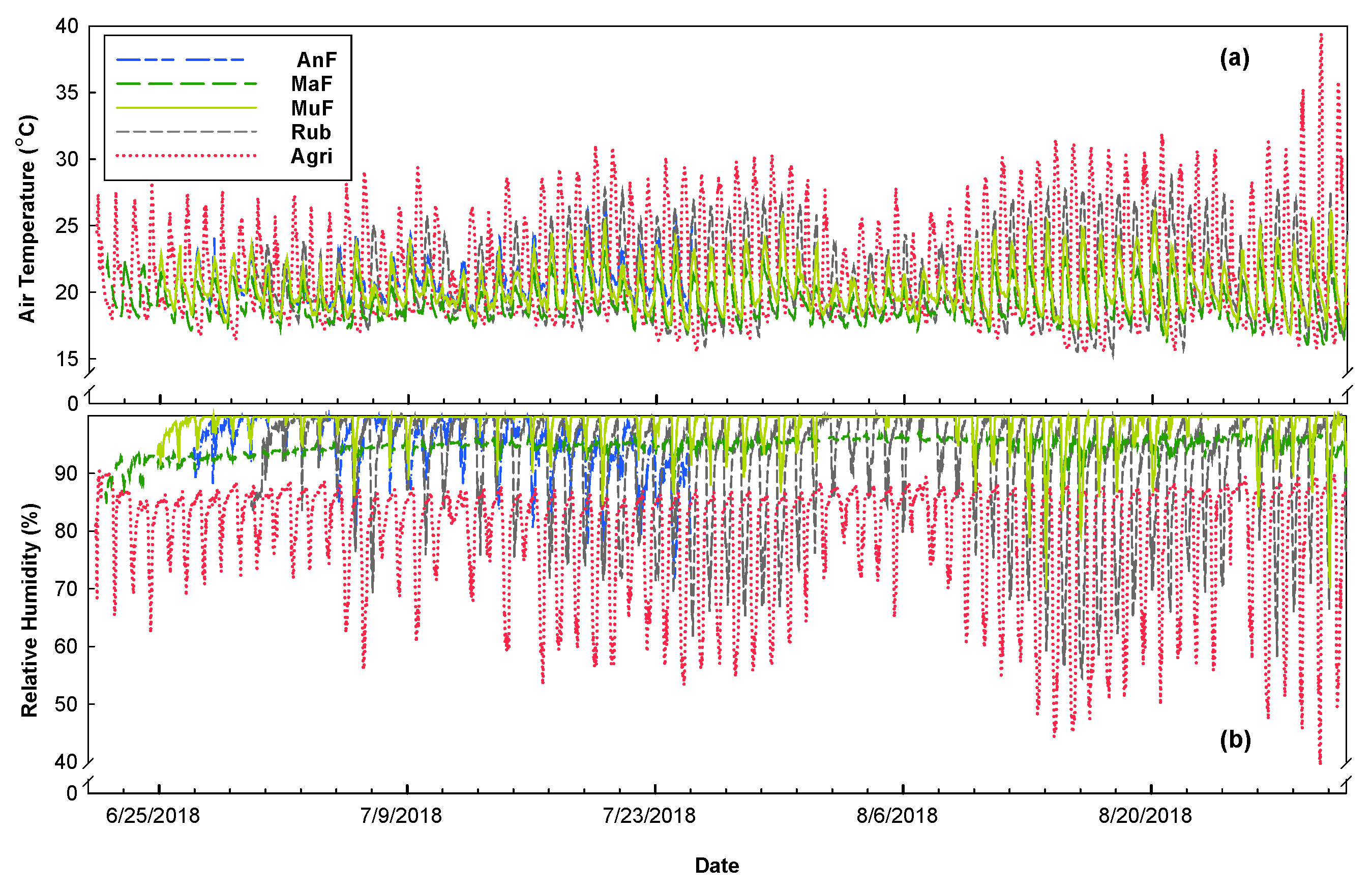

**Figure S2.** A modified USDA soil texture pyramid displaying the soil texture for the 5 study sites across a forest regeneration gradient: Angelo’s Forest (AnF), Mahua Falls Forest (MaF), Malungung Forest (MuF), Abandoned Rubber Plantation (Rub), and Abandoned Agriculture Field (Agri).

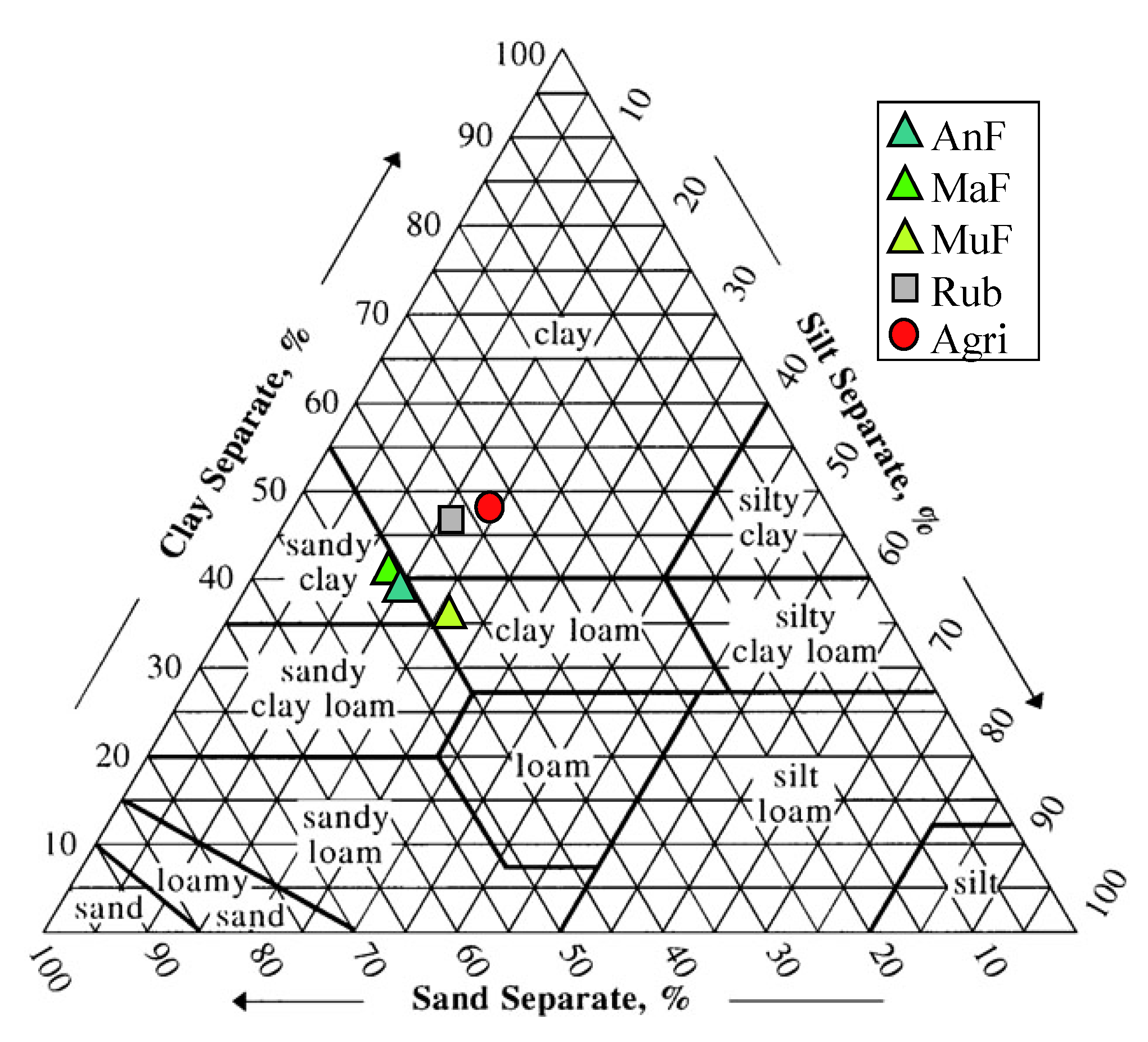

**Figure S3.** Relationship between recovery time and (a) total carbon (%), (b) total nitrogen (%), (c) litter decomposition rate (d^-1^), and (d) surrounding forest cover (%). Different sites including Angelo’s Forest (AnF), Mahua Falls Forest (MaF), Malungung Forest (MuF), Abandoned Rubber Plantation (Rub), and Abandoned Agriculture Field (Agri) are displayed using different colors and symbols. Smaller symbols in the background of each bivariate plot represent raw data points (n = 10), while larger symbols represent site means. Bars represent the standard error of mean and gray lines represent 95% confidence intervals. All statistical models are significant at p ≤ 0.05.

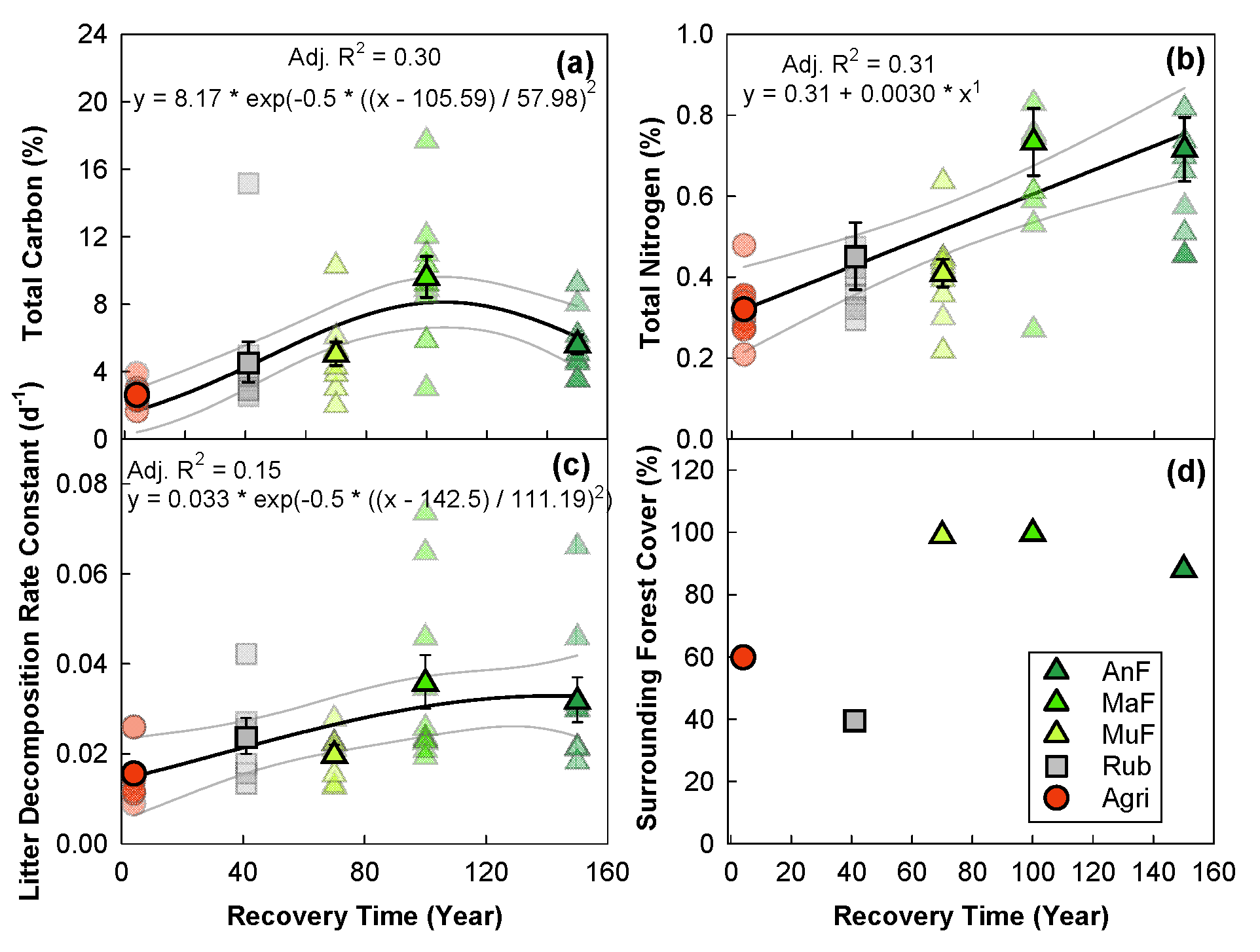

**Figure S4.** Relationship between litter stabilization factor, *S* (unitless) and litter decomposition rate, *k* (d^-1^) compared with global data [43]. Filled symbols represent site means: Angelo’s Forest (AnF), Mahua Falls Forest (MaF), Malungung Forest (MuF), Abandoned Rubber Plantation (Rubb), and Abandoned Agriculture Field (Agri); and numbered symbols represent a range of sites from global data: mangrove (1, 2); oceanic raised bog (3, 4); geothermal wet grassland (5, 6); semi-arid desert (7, 8); forest (9); wet forest (10); pasture (11); floating fen (12); lowland tropical forest (13); mixed forest (14); birch forest (15). Bars represent standard error. *Error bar absent due to overdispersion.

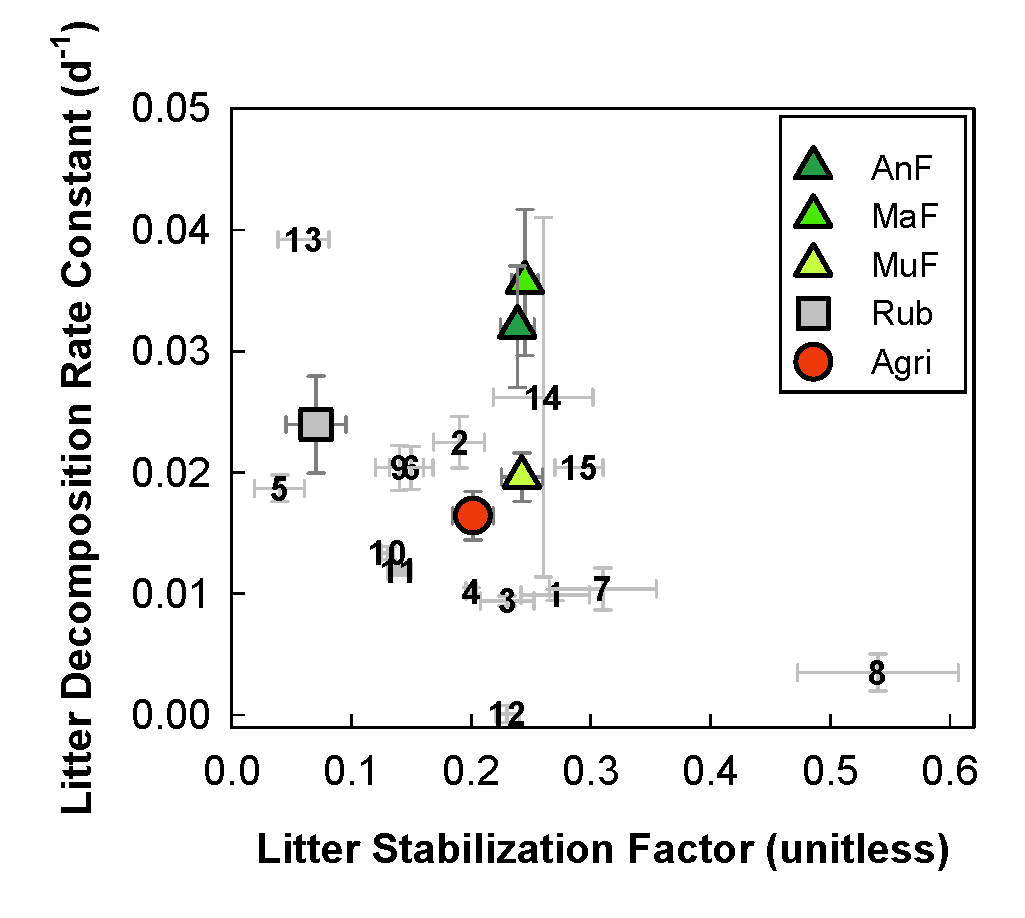

**Figure S5.** Relationship between surrounding forest cover (%) and daytime soil temperature (℃) across a forest regeneration gradient: Angelo’s Forest (AnF), Mahua Falls Forest (MaF), Malungung Forest (MuF), Abandoned Rubber Plantation (Rub), and Abandoned Agriculture Field (Agri). Smaller transparent symbols represent observed data points (n = 10), while solid symbols represent site means. Bars represent standard error of the mean, solid black line represents model mean, and gray lines represent 95% confidence intervals. Statistical model was run on observed data and significant at p < 0.001.

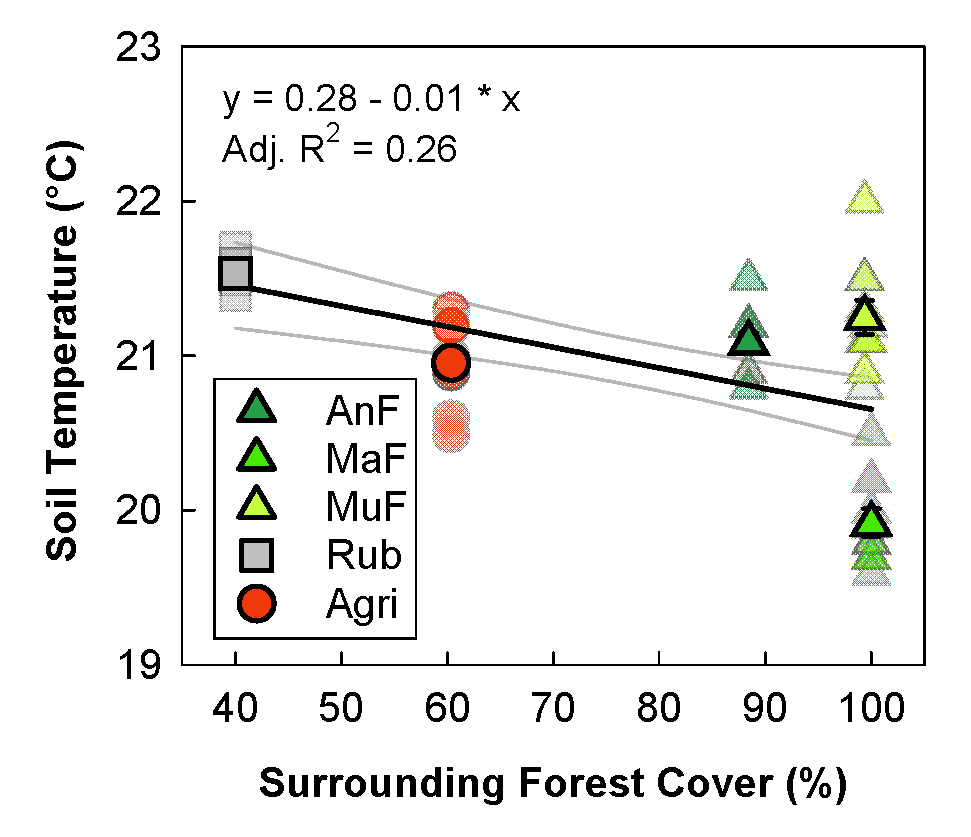

**Methods: Soil Microbial Analyses**

The 16s rRNA gene (bacteria) and ITS1 region (fungi) were amplified respectively. PCR primers 515, ITS1 and ITS2, along with HotStarTaq Master Mix Kit (Qiagen, USA) were used in 30 PCR cycles with the following steps: 94 ℃ for 3 min followed by 30 cycles of 94 ℃ for 30 s; 53 ℃ for 40 s and 72 ℃ for 1 min; and a final elongation step at 72 ℃ for 5 min*.* Following amplification, DNA fragments were sorted by molecular weight and concentration using 2% agarose gel*.*  The sorted DNA samples were then purified using an 1X AmpureXP beads and checked on Agilent High Sensitivity (HS) chip on Bioanalyzer 2100 and quantified on fluorimeter by Qubit dsDNA HS Assay kit (Brand from MR DNA). The library was loaded onto the Illumina Platform for clustering and sequencing. Paired-End sequencing allows the template fragments to be sequenced in both the forward and reverse directions. Sequencing was carried out on an illumina MiSeq following the manufacturer's guidelines and data were analyzed using the proprietary MR DNA analysis pipeline [35]. Sequences with < 150 bp or with ambiguous base calls were removed. Briefly, sequences were denoised, operational taxonomic units (OTUs) were assigned based on 97% similarity, and chimeras were removed. Singletons and OTUs appearing in only one sample were removed from OTU tables. Archaea and mitochondrial or chloroplasts OTUs were removed from the 16S data and non-fungi OTUs from ITS1 data. In total, 31 261 bacterial and 0 fungal chimeric were removed from further downstream analysis. Final OTUs were assigned using BLASTn; cloned sequences were searched against a database derived from RDP-II and NCBI (www.ncvi.nlm.nih.gov, <http://rdp.cme.msu.edu>). A total of 3 439 724 (68 794 ± 3 167 on average) high quality 16S sequences and 2 920 915 (58 418 ± 1 815) high quality ITS sequences for the 50 soil samples were analysed. The average of 1 274 ± 180 bacterial and 2 105 ± 59 fungal OTUs at 97% identity cutoff were obtained for 16S and ITS sequences.
